# Identification of Proteins Influencing CRISPR-Associated Transposases for Enhanced Genome Editing

**DOI:** 10.1101/2024.09.11.612086

**Authors:** Leo C.T. Song, Amanda T.P. Alker, Agnès Oromí-Bosch, Sophia E. Swartz, Jonathan N.V. Martinson, Jigyasa Arora, Abby M. Wang, Rachel Rovinsky, Sara J. Smith, Emily C. Pierce, Adam M. Deutschbauer, Jennifer A. Doudna, Brady F. Cress, Benjamin E. Rubin

## Abstract

CRISPR-Associated Transposases (CASTs) hold tremendous potential for microbial genome editing due to their ability to integrate large DNA cargos in a programmable and site-specific manner. However, the widespread application of CASTs has been hindered by their low efficiency in diverse, non-model bacteria. In an effort to address this shortcoming, we conducted the first genome-wide screen for host factors impacting *Vibrio cholerae* CAST (*Vch*CAST) activity and used the findings to increase *Vch*CAST editing efficiency. A genome-wide loss-of-function mutant library in *E. coli* was screened to identify 15 genes that impact type *Vch*CAST transposition. Of these, seven factors were validated to improve *Vch*CAST activity and two were found to be inhibitory. Informed by homologous recombination involved effectors, RecD and RecA, we tested the λ-Red recombineering system in our *Vch*CAST editing vectors, which increased its insertion meditated-editing efficiency by 25.7-fold in *E. coli* while maintaining high target specificity and similar insertion arrangements. Furthermore, λ-Red-enhanced *Vch*CAST achieved increased editing efficiency in the industrially important bacteria *Pseudomonas putida* and the emerging pathogen *Klebsiella michiganensis*. This study improves understanding of factors impacting *Vch*CAST activity and enhances its efficiency as a bacterial genome editor.

**GRAPHICAL ABSTRACT:** 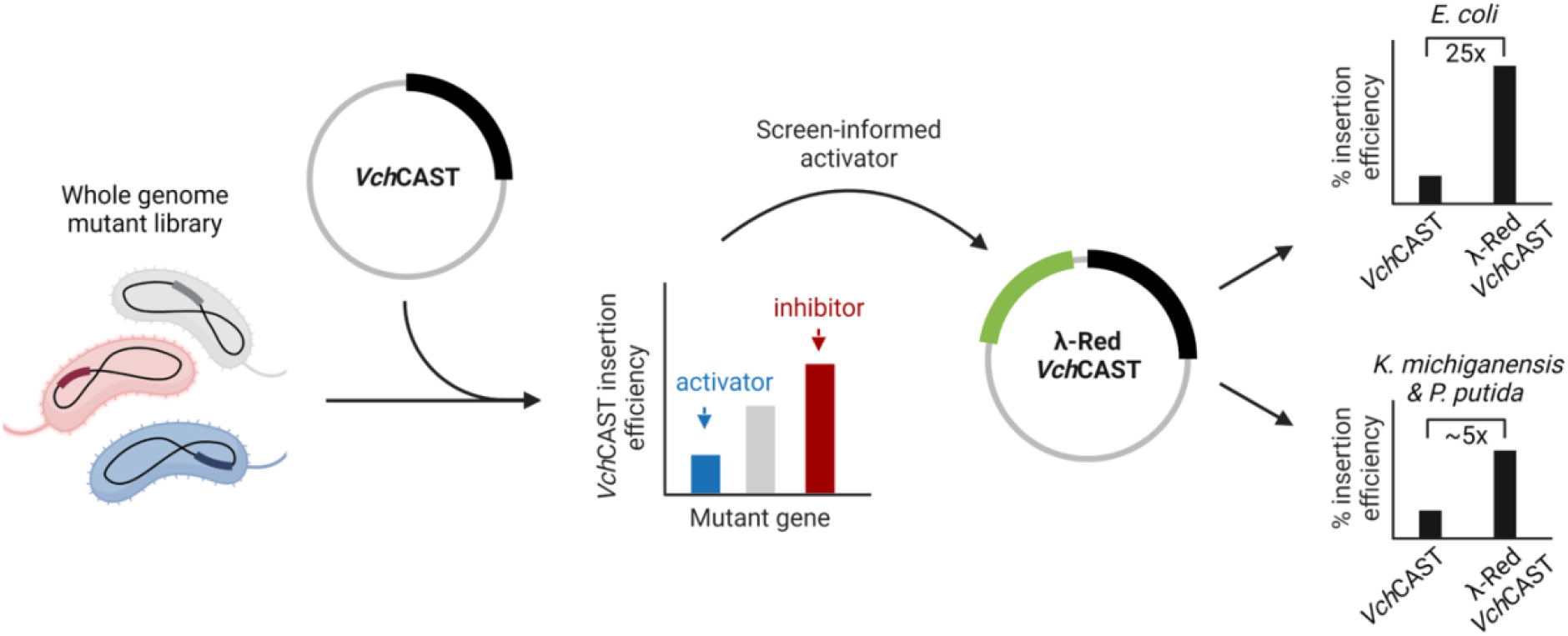

## INTRODUCTION

CRISPR-Associated Transposases (CASTs) represent a powerful new addition to the genome editing toolbox (1). Unlike traditional CRISPR-Cas systems that introduce targeted double-stranded breaks and rely on endogenous repair machinery to introduce edits, CASTs combine the RNA-guided targeting of CRISPR-Cas and Tn7-like transposition to make large programmable insertions (2). This unique mechanism circumvents the lethality often associated with double-stranded breaks in bacteria due to inefficient non-homologous end joining (3). For these reasons, a highly efficient and precise CAST, *Vibrio cholerae* type I-F CAST (*Vch*CAST) (Figure S1A), has emerged as a valuable system for bacterial editing (4), manipulating microbial communities (5), and even modifying human genomes (6).

Despite its established value as a genome editing tool, *Vch*CAST currently faces limitations in achieving efficient editing across a broad range of organisms. For example, in the industrially important species *Corynebacterium glutamicum, Vch*CAST achieved an editing efficiency between 0-0.027% (7). In human cells, the challenges are even more pronounced. Initial attempts to use *Vch*CAST in HEK293T cells resulted in editing efficiencies below 0.01% for genomic targets, with slightly higher rates of 0.1-1% observed only for episomal targets (6). In complex gut microbiome samples, *Vch*CAST has shown editing efficiencies as low as 0.001% (5), significantly limiting its utility for *in situ* microbiome engineering. These low efficiencies restrict *Vch*CAST’s potential for environmental, industrial, and therapeutic genome editing applications, highlighting the urgent need for optimization.

A significant barrier to broader and more efficient use of *Vch*CAST is the limited understanding of the host factors that enhance and restrict its function. For instance, the addition of ClpX was found to facilitate *Vch*CAST functionality in human HEK293T cells (6); however, ClpX has not been found necessary for its function in bacteria. Within bacteria, integration host factor (IHF) is the only protein complex known to affect *Vch*CAST function. A screen of variant *Vch*CAST transposon ends in *E. coli* showed that the presence of IHF is required for efficient transposition (8). However, a significant opportunity remains for systematic identification of activators and inhibitors for *Vch*CAST transposition.

In this study, we conducted a genome-wide screen to identify host factors that impact *Vch*CAST activity, resulting in the validation of seven activators and two inhibitors. Informed by the identification of homologous recombination-involved effectors, RecD and RecA, we explored whether incorporating the efficient phage-derived λ-Red recombination system could enhance *Vch*CAST editing performance. By integrating the λ-Red genes into our *Vch*CAST vectors, we achieved a 25.7-fold increase in editing efficiency. This strategy also facilitated more efficient integration in other industrially, environmentally, and medically relevant bacteria. Our research advances the understanding of CAST systems, revealing key regulatory factors and developing strategies to enhance editing efficiency. These improvements pave the way for broader application of CASTs across various host organisms, potentially enabling more efficient and versatile genome editing in both research and applied settings.

## MATERIALS AND METHODS

### RB-TnSeq screen for *E. coli* regulators of CAST integration

To systematically identify *E. coli* genes that contribute to *Vch*CAST-mediated integration, we conducted a genome-wide loss-of-function mutant screen and selected for *Vch*CAST edits after conjugation in the KEIO_ML9 *E. coli* RB-TnSeq library (9, 10). This approach allowed us to generate and select for *Vch*CAST edits across a diverse array of insertion mutants. The library comprises 152,018 uniquely barcoded single-gene transposon insertion mutants, covering 3,728 nonessential protein-coding genes out of a total of 4,146 in the *E. coli* genome (9). To account for biases introduced by the antibiotic resistance marker used, two different antibiotic selection cargos (P_Pmtl_-*gmR*, P_Pmtl_-*catP*) were utilized in the RB-TnSeq and conjugation efficiency assays, with a *mariner* transposase system serving as a control.

The KEIO_ML9 RB-TnSeq library was inoculated into LB containing kanamycin (25 µg/mL) and grown at 37ºC with shaking. Donor strains harboring the VcDART and *mariner* vectors respectively in *E. coli* WM3064 (11) (DAP auxotroph, *pir*^+^, RP4^+^) were grown at 37ºC with shaking in LB containing diaminopimelic acid (DAP; 0.3 mM) and gentamicin (screen 1; 50 µg/mL) or chloramphenicol (screen 2; 34 µg/mL). Three 10 OD*mL samples of the library overnight were washed and pelleted before being frozen at -80ºC as three technical replicate time zero (T=0) samples. After washing and resuspending in LB containing DAP, 1 OD*mL of donor was combined with 1 OD*mL of recipient in separate 1.5 mL Eppendorf tubes. Each combined donor-recipient sample was plated onto a plain LB agar petri plate topped with a MF-Millipore™ Membrane Filter for conjugation. After 6 hours of conjugation at 30ºC, cells from two conjugation plates were scraped into 20 mL of LB as a single technical replicate (six conjugation plates for three technical replicates for each donor-recipient combination). 100 uL of the resuspended cells were set aside for 10-fold serial dilution and spot plating. The remaining cells from each technical replicate were plated onto selective BioAssay dishes of LB agar containing kanamycin and gentamicin or chloramphenicol. The three technical replicates of the library not introduced to any donor were plated on nonselective BioAssay dishes of LB agar with kanamycin. The BioAssay dishes were incubated at 30ºC for 12 hours. Cells on BioAssay dishes were then scraped and resuspended in 20 mL of LB. A two mL aliquot of the resuspended cells was taken for genomic DNA extraction via the QIAGEN DNeasy PowerSoil Pro Kit. Barcodes were amplified with the BarSeq_v3 primers following the BarSeq PCR protocol (9). Amplicons were pooled into a synthetic amplicon library and submitted for sequencing by Illumina NovaSeq PE150.

BarSeq sequencing reads were processed using the FEBA pipeline described previously (9). In this pipeline, the final gene fitness value was calculated by averaging the fitness scores of all strains with independent transposon insertions located in the central region of the gene, excluding those near the beginning or end. The pipeline-generated fitness scores (fit_logratios_good) were further analyzed using R (version 4.4.0) with the tidyverse package (version 2.0.0). Within an experimental trial, treatment replicates were averaged and then the difference between the *Vch*CAST fitness scores and the *mariner* fitness scores was calculated. To identify genes consistently displaying strong differential effects, we focused on those with an absolute fitness difference greater than one in both experimental trials. Positive fitness values generated by the FEBA pipeline indicate a putative inhibitor of *Vch*CAST, as mutating the gene increases the fitness of the strain. Conversely, negative fitness values indicate putative activators of *Vch*CAST function.

### Validating activators and inhibitors with Keio *E. coli* mutants

We next validated the involvement of putative host factors identified in the RB-TnSeq screen by performing *Vch*CAST editing in *E. coli* Keio collection deletion mutants corresponding to the genes with the largest fitness values (absolute gene fitness > 1) and possessing molecular functions of interest as detailed on the Gene Ontology (GO) database (12). To confirm the Keio strains, primers were designed to amplify the kanamycin resistance gene insertion within each target locus of the expected Keio mutants (13) (Table S1). The *ΔyicI* mutant was selected as the negative control as it is a context-neutral genomic locus with no documented adverse fitness effects (14).

To assess how the presence or absence of individual *E. coli* genes affect *Vch*CAST integration into the target genome, we implemented a conjugation-based editing efficiency assay (4) (Figure S1B). Briefly, donor and recipient strains were grown for 16 hours, washed, resuspended in LB containing DAP, combined in a 1:1 ratio (0.1 OD*mL each), spotted on LB agar within a 24 well block, and allowed to conjugate for 6 hours at 30ºC. Afterward, spots were resuspended in one mL LB media. Ten-fold serial dilutions were performed with resuspended cells, spotted onto LB agar and LB agar with antibiotics, and incubated overnight at 30ºC. Serial dilution spot plates were left to grow until individual colonies formed. Finally, the plates were imaged and cell colonies were counted to compute editing efficiency (4, 5), with transconjugants reflecting successful conjugation and insertion of selective cargo. Fold change in editing efficiency was computed by normalizing the editing efficiency of candidate regulator hits of interest to that of the *ΔyicI* control. Three biological replicates were used for each experimental condition.

### Construction and testing λ-Red *Vch*CAST in *E. coli*

The DNA of λ-Red and Jungle Express (pJEx) were purchased as gblock gene fragments (IDT) and added to *Vch*CAST vectors using Gibson and Golden Gate assemblies. For testing the effect of the phage-derived homologous recombination system on editing efficiency we cloned λ-Red onto a *Vch*CAST backbone just after the origin of transfer, ensuring quick transcription in the recipient cell during conjugation. The tightly regulated and strong, inducible Jungle Express promoter (pJEx) was used to minimize leaky λ-Red expression and toxicity of the vector. pJEx is inducible with crystal violet (CV) and transcriptional control is robust in Pseudomonadota (15).

Plasmids, strains, synthesized DNA, and oligonucleotides (IDT, Coralville, USA) used in the study are listed in supplemental materials (Table S1-3). *Vch*CAST vectors were assembled through multipart Golden Gate cloning. High-fidelity PCRs were performed with Q5 Hot Start High-Fidelity DNA polymerase (NEB). Golden Gate assembly enzymes (e.g. BsmBI-V2, BbsI, BsaI-HFV2, and T4 ligase) were ordered from NEB and used with the reported buffers following previously reported protocols (5). *Vch*CAST assemblies were electroporated into electrocompetent *E. coli* EC100D*pir*+ cells (LGC Biosearch). Clones were screened by colony PCR (cPCR) with 2x GoTaq Green Mastermix (Promega) and plasmids were isolated with a QIAprep Spin Miniprep Kit (Qiagen). Guide assemblies were electroporated into *E. coli* WM3064 *pir*+ and grown on the appropriate antibiotics plus DAP. All vectors were confirmed with whole-plasmid sequencing (Plasmidsaurus Labs).

Crystal violet induction was tested in *E. coli* to determine the concentration that produces the highest insertion efficiency (Figure S2). Conjugations were performed on agar plates containing the crystal violet at the optimal induction concentration (0.01 µM) for 6 hours before resuspension and selection. To account for variability in editing efficiency between biological replicates, λ-Red *Vch*CAST treatments were paired and normalized to the negative *Vch*CAST control. Each experiment was repeated with three biological replicates.

### Testing λ-Red *Vch*CAST in *P. putida* and *K. michiganensis*

We first performed a quantitative assay in candidate strains to determine the frequency of mutants resistant to the antibiotics streptomycin/spectinomycin/carbenicillin (100 µg/mL and 400 µg/mL), chloramphenicol (34 µg/mL and 68 µg/mL), kanamycin (25 µg/mL, 50 µg/mL, 100 µg/mL and 200 µg/mL), and gentamicin (10 µg/mL, 20 µg/mL, and 40 µg/mL). Overnight cultures of each species were grown in LB at 30ºC, 10x serially diluted, and spotted onto LB agar without antibiotics (control) and onto each of the antibiotic concentrations. Colonies were counted after 16-40 hours of growth, and the antibiotic concentration exhibiting minimal or no detectable growth was chosen as a selection marker for genome editing experiments (Figure S5A-B). Based on these results, we constructed a *Vch*CAST vector containing the P_pmtl_ promoter driving the kanamycin resistance gene for selection in *P. putida* and *K. michiganensis*.

We identified safe sites and designed guides in *P. putida* (GCF_000007565.2) following previously reported methods (4, 5) and used a previously tested safe site guide in *K.michiganensis (5)*. Briefly, intergenic regions between converging genes with a distance of 300-600 nucleotides were selected (Figure S5D). Candidate safe sites were excluded under any of the following circumstances: the region was located within or adjacent to predicted mobile genetic elements (MGEs), the region was flanked by essential genes (inspected using BioCyc), or the region contained non-coding RNA (ncRNA) features (inspected using Rfam). Within the selected safe site regions, protospacer adjacent motifs (PAMs) with the sequence 5’-CN-3’ were identified. For each PAM, 32 nucleotides were added to generate a list of potential guides, which were chosen based on having a GC content of 40-60%, and ensuring that the insertion loci, approximately 49 bp downstream of the guide, remained within the intergenic region and that it would not accidentally disrupt a terminator sequence. Three guides per safe site were selected, and their off-target potential was assessed using a local BLASTn search (-dust no - word_size 4). Guides with off-target hits exhibiting the highest e-values and with minimal complementarity to off-targets in the seed region (first ∼10 nt) were prioritized. Two guides targeting different safe sites were cloned and tested (Figure S5C).

Conjugation experiments in *P. putida* and *K. michiganensis*, were performed following the protocol described for *E. coli* with some modifications. Recipient and donor cells were grown 16 hours, at 30 ºC and 37 ºC, respectively. Conjugations were done for 20 hours at 30 ºC with crystal violet induction concentrations of 0.5 µM and 1 µM used for *K. michiganensis* and *P. putida*, respectively (Figure S5E-F). Transconjugants were selected on LB plates containing kanamycin (50 µg/mL). Drip plates of 10 µL for each serial dilution were plated to increase sensitivity and reduce technical error. Three biological replicates were performed across different days.

### Insertion analysis in *E. coli, P. putida* and *K. michiganensis*

Insertion analysis was preliminarily performed by cPCR on transconjugants following selection in all strains. Insertion orientation primers were designed to identify clones with right-left (T-RL) and left-right (T-LR) simple insertion, cointegration, and no integration outcomes (4) (Figure S3A, S3C). Cointegration oligos were designed to amplify the junction between the *Vch*CAST vector backbone and the resistance marker in the cargo. Simple insert oligos were designed to amplify off of the flanking genomic DNA to capture the full *Vch*CAST cargo insert. Another set of oligos were designed to distinguish between T-RL versus T-LR simple insert products.

Additional screening for insertion cointegration versus simple insertion was performed for *E. coli* by patch plating of transconjugants on LB agar plates containing 100 μg/mL carbenicillin, the *Vch*CAST vector backbone resistance marker. Carbenicillin resistance in *P. putida* and *K. michiganensis* did not permit cointegrate screening by double selection and required insert orientation analysis after selection either by cPCR or WGS.

Insertion products from all recipient strains in this study (*E. coli, P. putida* and *K. michiganensis*) were further assayed for off-targets by whole-genome sequencing. To address colony heterogeneity, which has been observed previously (4), about a thousand colonies of transconjugant cells transformed with λ-Red *Vch*CAST, were scraped from selection plates and resuspended. For each species investigated, resuspensions from three biological replicates were pooled together and replated on the appropriate dilution to yield thousands of single colonies. Colonies were scraped from selection plates and resuspended in a volume of LB media equivalent to OD_600_=3-4. An aliquot of this resuspension was used for high-molecular-weight genomic DNA (gDNA) extraction using the MasterPure™ Complete DNA and RNA Purification Kit (Biosearch Technologies). Genomic DNA samples were submitted for Oxford Nanopore long-read sequencing to Plasmidsaurus Labs.

The bioinformatics analysis was performed using custom scripts. The demultiplexed raw reads were filtered using nanofilt python package (v 2.8.0) to select reads with a Phred quality score of 20 (>Q20) and a minimum length of 150 base pairs (bp) (16). To identify the inserted cargo, the reads were aligned to the last 71-bp of the transposon right end, which consists of three 20-bp TnsB binding sites and a 8-bp terminal repeat. The alignment was performed using minimap2 software (17) and reads mapping to less than 70% of the right end of cargo were filtered out.

For downstream analysis, a pipeline developed by Vo et al. (2021) was employed with some modifications (18). The reads containing the mapped region were extracted and aligned with the complete reference and plasmid genomes. The reads were classified as genomic, plasmid, or both based on whether more than 500 bp was mapped to the respective genomes.

The genomic coordinates of the mapped region were recorded to generate the genome-wide histograms of the integration location. The mapped region was categorized as on-target if it fell within the 100-bp window of the 3’end of the target site.

### Bioinformatic homology search for regulators

We examined the phylogenetic distribution of 11 genes across 92 bacterial phyla, which included the nine identified and validated by the RB-TnSeq screen plus *ihfA* and *recA*. To simplify viewing, only phyla with more than 10 members in the AnnoTree database (v beta, based on GTDB R214.0 (19)), were included in this analysis. Briefly, the genes in the AnnoTree database were annotated using PFAM v27.0 (20), TIGRFAM v15.0 (21), and KEGG orthology identifiers assigned to the UniRef100 database (22). The KEGG or PFAM annotation assigned to individual genes was used to extract homologs from 80789 representative genomes from GTDB R214.0 (23). Genes such as *ihfB, recD*, and *umuD* are found in close association with their respective enzymatic complex subunits, *ihfA, recBC*, and *umuC* (24–26). To maintain consistency, the homologs of these co-occurring genes were extracted using the same methods as for the genes of primary interest, and only genomes containing both sets of genes in the same contig were retained for further analysis. The gene *yneK* lacks a PFAM or KEGG annotation. To address this, protein sequences from the 80,789 representative genomes in the GTDB database (23) were downloaded and subjected to BLASTp analysis against the *E. coli yneK* sequence. Proteins with greater than 50% sequence similarity and a significant FASTA hit (E ≤, 1e-5) were retained. To further validate our findings, a BLASTp search was conducted against a UniProt database subset containing sequences sharing at least 90% similarity to E. coli yneK. Both approaches yielded comparable results. While the yneK gene is prevalent among closely related *E. coli* and *Shigella* strains, only a limited number of these strains are present in the 80789 representative genomes in the GTDB database. The *cspC* gene is a member of the bacterial Cold Shock Protein (CSP) family (27), which can contain multiple paralogs with high sequence similarity (>70%; (28). To accurately differentiate these paralogs, cspC proteins identified by AnnoTree were re-analyzed using eggNOG-mapper v5.0 (29). This tool leverages evolutionary relationships to distinguish between proteins within the CSP family excluding non-*cspC* family members. Finally, gene presence per phyla was visualized in iTOL v6 (30) with the publicly available GTDB R214.0 bacterial tree inferred from the concatenation of 120 proteins (31, 32).

## RESULTS

### A whole genome mutant screen and validation to identify *Vch*CAST regulators

To perform a genome-wide survey of genes involved with *Vch*CAST function, we screened a library of loss-of-function mutants for the ability of *Vch*CAST to integrate into a safe site. The screen was conducted with a preexisting *E. coli* RB-TnSeq transposon mutant library (9).

*Vch*CAST was introduced to the library via conjugation, and the inserted cargo was selected for, ensuring all members of the library contained both the *Vch*CAST and the background loss-of-function mutation (Figure 1A). The efficiency of *Vch*CAST insertion into each mutant was then determined by assessing the abundance of each insertion-containing mutant by sequencing their unique barcode. We identified eleven genes as putative activators and four as putative inhibitors, based on their consistent phenotypes (absolute gene fitness > 1) in two screens conducted in parallel (Figure 1B). Notably, the beta subunit of integration host factor (*ihfB*) emerged as the top activator from the screen (Figure 1B). The other subunit, *ihfA*, did not have a high enough abundance in the T=0 starting library to be considered in the analysis pipeline. This finding aligns with previous research, as IHF is the only currently known activator of *Vch*CAST insertion (8), lending credibility to the screen’s results.

**Figure 1.**
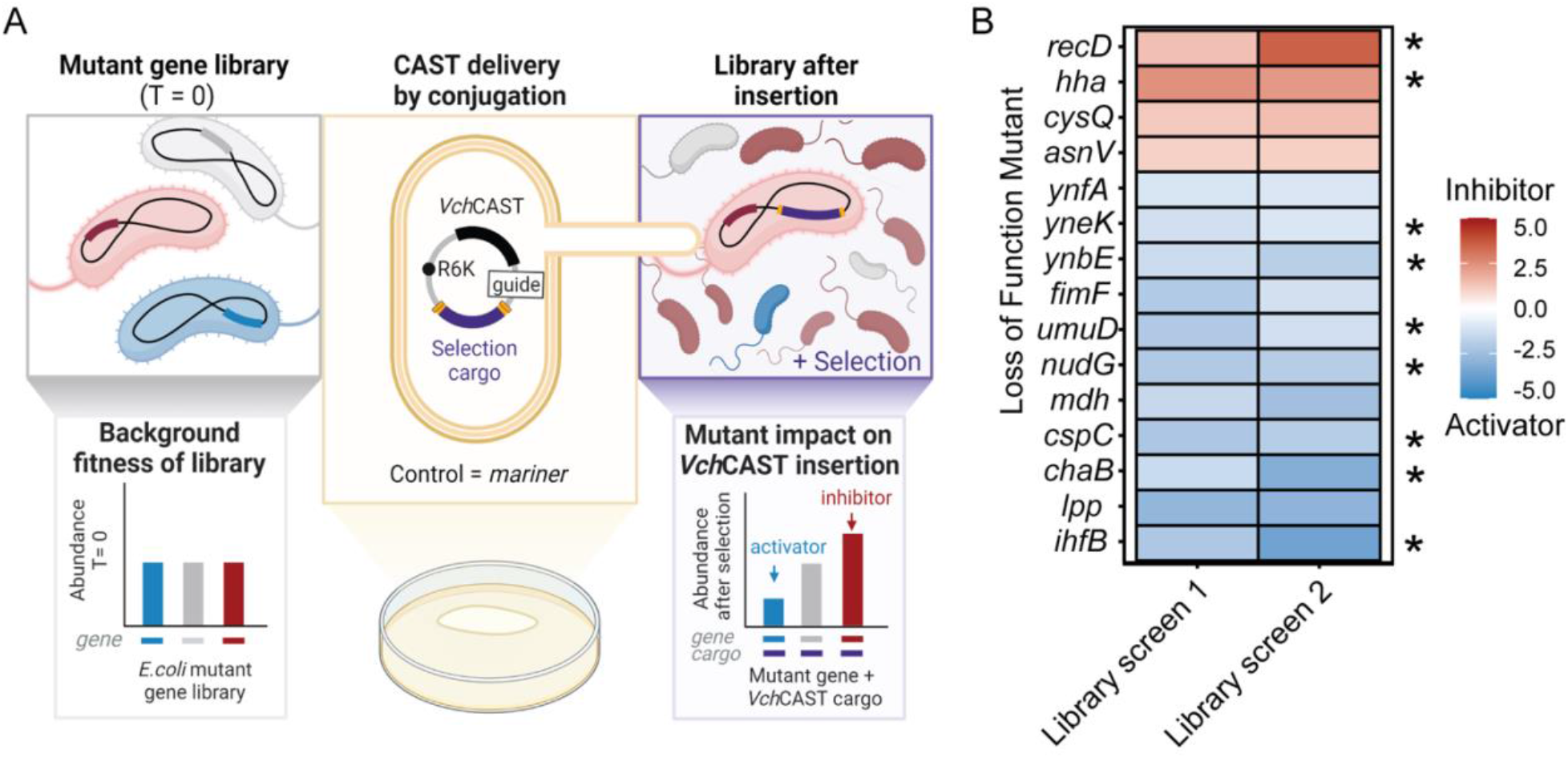
Genome-wide screen identifies putative inhibitors and activators of VchCAST integration. (A) Schematic of the RB-TnSeq screen to identify *E. coli* genes affecting *Vch*CAST integration efficiency. *Created in BioRender. Alker, A. (2024)* BioRender.com/j90j989 (B) Fitness scores of *E. coli* genes from two independent RB-TnSeq screens with fitness scores >1 or <-1 in both screens. Library screens 1 & 2 refer to repeated screens with gentamicin and chloramphenicol selection cargo, respectively. Stars denote candidate mutants that were experimentally validated.

We validated hits with the largest absolute fitness values from the pooled RB-TnSeq screens by testing *Vch*CAST editing in clonal gene deletion mutants. To prioritize candidates most likely to have direct interaction with *Vch*CAST, we filtered for those possessing DNA-interacting, RNA-interacting, and protein-interacting functions as well as hypothetical proteins, as per the Gene Ontology (GO) database (Table S4). We then performed conjugation assays to quantify the relative editing efficiency of *Vch*CAST in Keio deletion mutants of each of the nine filtered hits (Figure 2A). Editing efficiencies were compared to *ΔyicI*, a deletion mutant with no documented adverse fitness effects (14). Overall, mutants of putative activators (*ihfB, cspC, ynbE, umuD, chaB, nudG, yneK*) exhibited reduced *Vch*CAST editing efficiency by 7.7 ± 21.7% to 99.9 ± 0.1% compared to the control. Knockouts of putative inhibitors (*hha, recD*) exhibited increased editing efficiency (4.0 ± 1.6-fold and 21.8 ± 7.2-fold) in comparison to the *ΔyicI* control (Figure 2A). The results from the whole-genome screen and the individual mutant validation were largely consistent, with *recD* and *ihfB* showing up in both experiments as the strongest inhibitor and activator, respectively. Taken together, the validation results support the involvement of these host factors in *Vch*CAST’s genomic integration.

**Figure 2.**
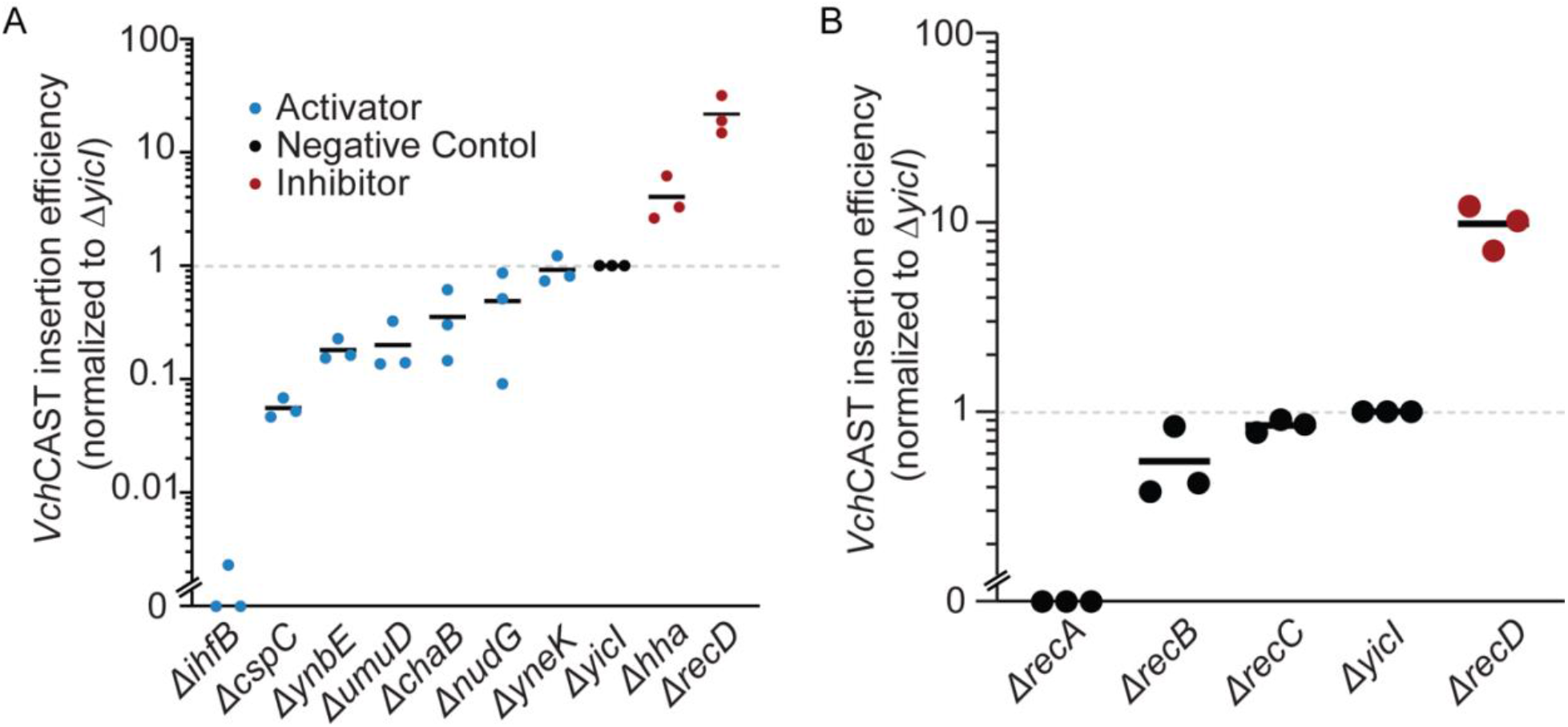
Validation of putative VchCAST activators and inhibitors in single deletion mutants. (A) Relative *Vch*CAST editing efficiency in *E. coli* Keio collection mutants of candidate regulators identified from the RB-TnSeq screen. The editing efficiencies are normalized to the Δ*yicI* control strain. Data values at zero represent samples with no viable colonies above the detection limit. (B) Relative *Vch*CAST editing efficiency in *E. coli* Keio collection mutants of RecBCD complex components and downstream homologous recombination effector RecA normalized to the Δ*yicI* negative control strain. Data values at zero represent samples with no viable colonies above the detection limit.

We were particularly compelled by the strong inhibitor classification of *recD* in *Vch*CAST insertion revealed by the screen and validation, and sought to explore the *rec* genes further. RecD*’s* role in the RecBCD complex leads to the inhibition of homologous recombination (33), suggesting a beneficial role of factors involved in DNA repair during *Vch*CAST integration. To test this hypothesized role, we delivered *Vch*CAST via conjugation into Keio knock-out mutants of select, key *rec* genes (*recABCD*) and quantified editing efficiency normalized to *ΔyicI* (Figure 2B). Consistent with the screen results, the putative inhibitor *ΔrecD* showed a 9.8 ± 2.6-fold increase in editing efficiency compared to the neutral fitness mutant control. Knockouts of *recB* and *recC* slightly decreased insertion efficiency relative to the control (45.3 ± 25.5% and 15.0 ± 6.6%, respectively). *ΔrecA*, which did not have a high enough abundance in the T=0 starting library to be considered in the RB-TnSeq screen, had no viable transconjugant colonies above the limit of detection. The results of this experiment identify *recA* as an activator and support the role of homologous recombination in promoting *Vch*CAST integration.

### Leveraging λ-Red to improve *Vch*CAST editing efficiency in *E. coli*

Considering the role of the RecBCD complex and potentially homologous recombination more broadly in *Vch*CAST integration, we hypothesized that introducing higher-efficiency recombination machinery may improve editing outcomes. To this end, the bacteriophage λ-Red genes (*exo, beta*, and *gam*) were cloned into a *Vch*CAST plasmid (R6K, P_Pmtl_-*catP* cargo) (Figure 3A). We optimized the induction level of λ-Red by crystal violet (Figure S2) for editing in *E. coli*. The induced λ-Red *Vch*CAST treatment significantly increased editing efficiency by 25.7 ± 0.6 fold in BW25113 *E. coli* compared to the *Vch*CAST control (Figure 3B). The non-targeting (NT) control did not yield colonies within the level of detection for our assay. Insertion analysis of *E. coli* transconjugants by cPCR (n = 30 colonies) showed the same simple insert and co-integration rates between λ-Red *Vch*CAST (86.7% simple insert, 13.3% co-integrate) and *Vch*CAST (86.7% simple insert, 13.3% co-integrate) (Figure S3B), corresponding to previously reported rates (34). Orientation of simple inserts were also similar between λ-Red *Vch*CAST (92.3% T-RL, 7.7% T-LR), *Vch*CAST (96.2% T-RL, 3.9% T-LR), and previously reported values (34). WGS analysis confirmed that 100% of λ-Red *Vch*CAST insertions were on-target in *E. coli* (Figure 3C). These data support λ-Red’s positive impact on *Vch*CAST editing efficiency without impacting insertion outcomes.

**Figure 3.**
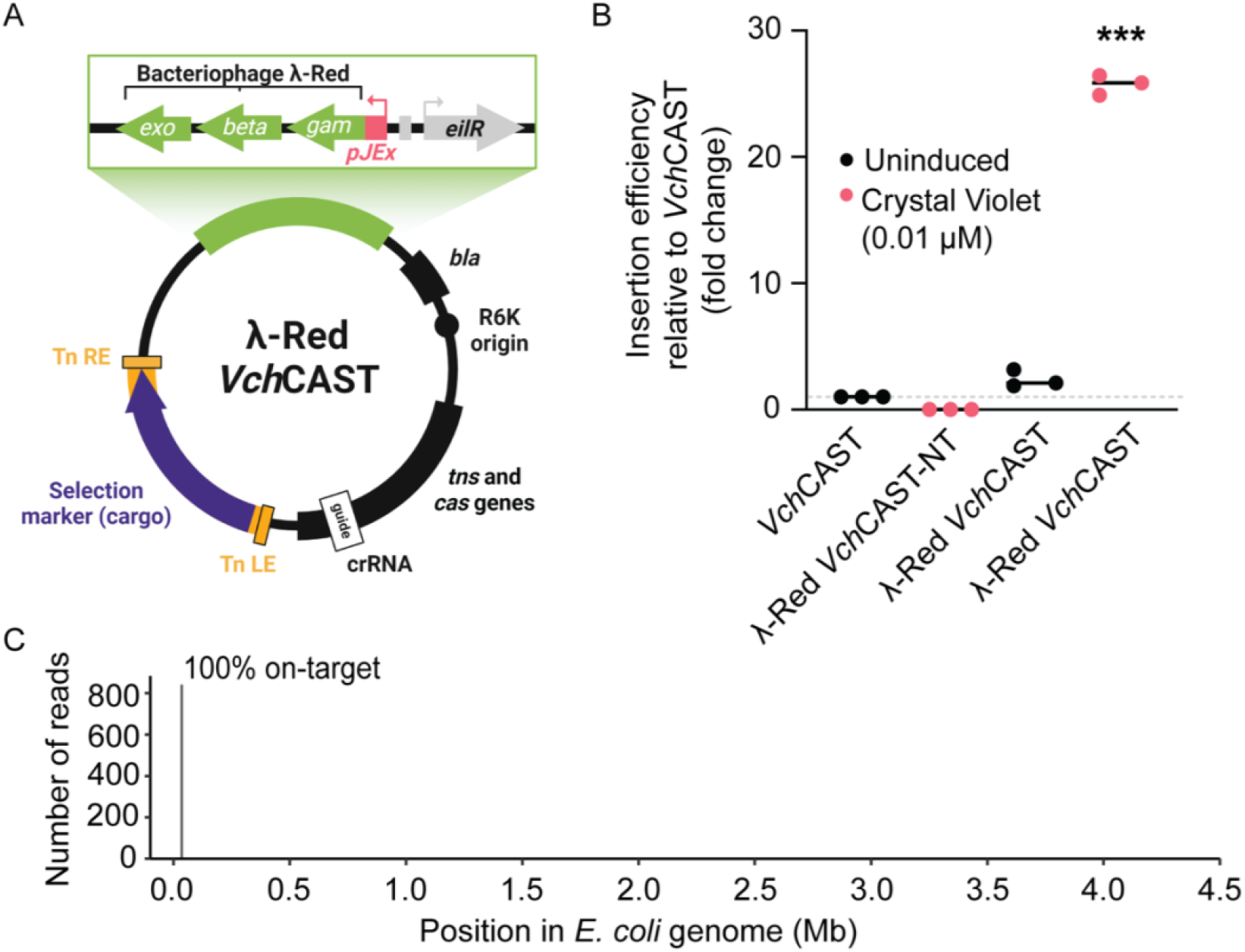
λ-Red recombineering system improves VchCAST editing efficiency in E. coli. (A) Schematic of the λ-Red *Vch*CAST vector design. Created in BioRender. Alker, A. (2024) BioRender.com/o37n467 (B) Comparison of editing efficiency between *Vch*CAST and λ-Red *Vch*CAST in *E. coli* BW25113. The editing efficiency of λ-Red *Vch*CAST as well as the non-targeting (NT) biological replicates were normalized to the paired *Vch*CAST replicates. *** p = 0.0003 (One sample two-sided t-test compared to hypothetical mean (1) of *Vch*CAST). (C) On-target insertion frequency of *E. coli* transconjugants via whole genome sequencing.

We next tested which of the λ-Red genes on *Vch*CAST were necessary for the observed increase in editing efficiency. Constructs containing inducible *exo-beta, gam* only, and the full λ-Red operon on the backbone of the CAST editing vector were tested. Exo and Beta promote homologous recombination through their exonuclease and single-stranded binding activity respectively, while Gam inhibits RecBCD (35, 36). We found that while the induced exo-beta construct does improve editing efficiency relative to the *Vch*CAST*-*only control by 6.9 ± 1.8-fold, it does not improve editing to the extent of all three lambda genes (28.8 ± 18.5-fold) (Figure S4). The presence of *gam* alone on the *Vch*CAST vector did not affect editing efficiency (Figure S4). Taken together, these results suggest that the complete λ-Red system is necessary for maximizing the efficiency of *Vch*CAST integration.

### λ-Red-assisted *Vch*CAST editing in additional gram-negative bacteria

Motivated by the successful implementation of λ-Red to increase *Vch*CAST-directed editing efficiency in *E. coli*, we evaluated the impact of λ-Red on *Vch*CAST-mediated DNA integration in other gram-negative bacteria. We selected *P. putida* KT2440, a relevant species for applications in industrial biotechnology and bioremediation (37), and *K. michiganensis* M5a1, a plant-associated nitrogen-fixing strain and close relative to an important human pathogen (38, 39).

The *Vch*CAST and λ-Red *Vch*CAST vectors were conjugated into both species and queried for their effect on insertion efficiency. In both species, expression of λ-Red from the *Vch*CAST vector resulted in increased insertion efficiency compared to *Vch*CAST alone (Figure 4A-B). In *P. putida*, we observed an increase with (3.6 ± 1.1 fold-increase) and without (4.7 ± 1.6 fold-increase) the inducer (1 µM CV) suggesting that leaky expression of λ-Red from the pJEx promoter in this strain is sufficient to enhance editing efficiency (Figure 4A). In *K. michiganensis*, upon induction with 0.5 µM CV, there is a 7.7 ± 7.0-fold increase in percent editing efficiency (Figure 4B).

**Figure 4.**
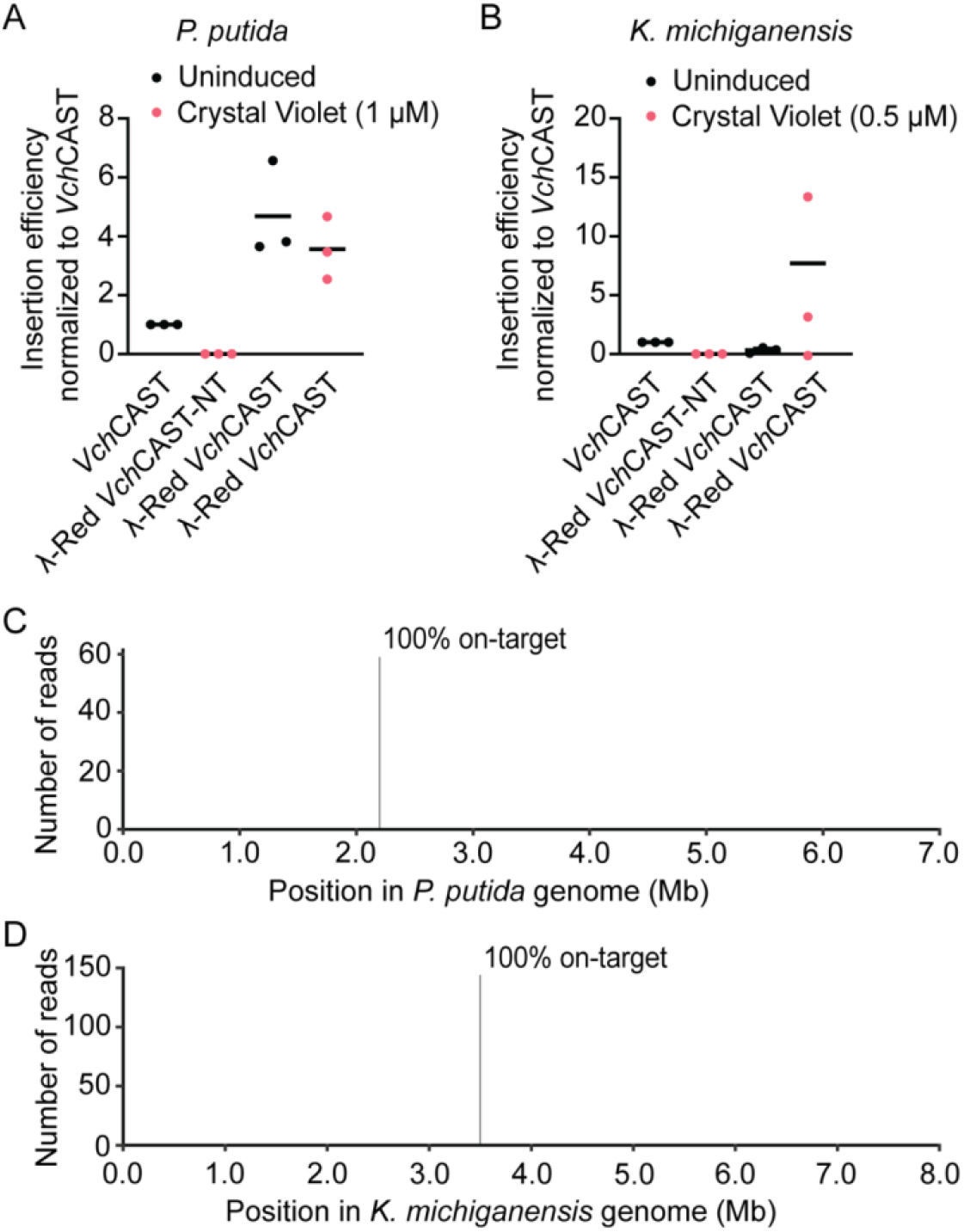
λ-Red improves VchCAST editing efficiency in P. putida and K. michiganensis. Editing efficiency of *Vch*CAST and λ-Red *Vch*CAST in (A) *P. putida* and (B) *K. michiganensis* with (1 µM and 0.5µM, respectively) and without CV induction. The editing efficiency of λ-Red *Vch*CAST as well as the non-targeting (NT) biological replicates were normalized to the paired *Vch*CAST replicates. (C) On-target insertion frequency of *P. putida* transconjugants via whole genome sequencing. (D) On-target insertion frequency of *K. michiganensis* via whole genome sequencing.

We analyzed the *Vch*CAST and λ-Red *Vch*CAST insertion products by WGS analysis in both *P. putida* and *K. michiganensis*. In both bacteria, RNA-guided DNA insertions were 100% on-target for the colonies assayed (Figure 4C-D). Altogether, these results highlight the potential of using λ-Red *Vch*CAST to enhance on-target *Vch*CAST editing efficiency for precise genome engineering in diverse gram-negative bacteria.

### Distribution of *Vch*CAST inhibitors and activators across diverse bacteria

To understand the broader relevance of our findings, we investigated the phylogenetic distribution of *Vch*CAST activators, inhibitors and associated genes across the bacterial tree of life. We analyzed the presence of our seven validated RB-TnSeq activators, two inhibitors, as well as *ihfA* (8) and *recA* (shown to be a strong activator in this study) across all bacterial phyla with more than 10 members in the AnnoTree database. Our analysis revealed diverse conservation patterns among these genes. RecA has near-universal presence across the microbial tree of life and was observed in over 90% of species in most phyla. In contrast, other genes identified from the screen such as *yneK, hha, yneB*, and *chaB* are absent across most phyla beyond *E. coli* containing Pseudomonadota and present only in small portions of the phyla where they exist. This landscape of *Vch*CAST activators and inhibitors should provide insight into where *Vch*CAST may be used effectively and what modifications may be necessary to make it function in organisms where it currently does not.

## DISCUSSION

In this study, we conducted a genome-wide mutant screen to identify genes in *E. coli* that influence the efficiency of *Vch*CAST, a promising tool for precise DNA insertion of large cargos in bacteria and eukaryotes. We screened for and then validated nine candidate genes, with eight being novel to this study, that either positively or negatively affect *Vch*CAST insertion activity when disrupted (Figure 1B, 2A). Notably, our results highlighted a role for the RecBCD complex in CAST integration, with the disruption of *recD* increasing editing efficiency, while a *recA* deletion strongly decreased it (Figure 2B). Building on these insights, we leveraged the bacteriophage λ-Red genes (*exo, beta*, and *gam*) to enhance CAST insertion efficiency (Figure 3A). By optimizing the expression of these genes, we achieved improved editing efficiency not only in *E. coli* (Figure 3B), but also in *P. putida* (Figure 4A), an industrial model strain (37), and *K. michiganensis* (Figure 4B), a plant-associated nitrogen-fixing strain and close pathogen relative (39). This work provides a comprehensive survey of host factors influencing *Vch*CAST integration and presents an approach to enhance its efficiency across various bacterial species.

Beyond the homologous recombination-associated RecD, we discovered eight other activators and inhibitors of the CAST complex that may further contribute to understanding *Vch*CAST transposition (Figure 2A). Putative activator UmuD, through its role in inhibiting DNA polymerase III from binding to ssDNA (40), may protect the integration site from premature replication fork collision. Cold shock protein C (CspC), known to bind to single-stranded nucleotides (41), could stabilize the RNA guide or the single-stranded gap of the post-strand transfer intermediate. Interestingly, ClpX, a sequence-specific AAA+ ATPase with protein unfolding capabilities that facilitated more efficient editing in human cells (6), was not identified in our screen. It is possible that in *E. coli*, one or more redundant proteins can perform ClpX’s activity. The new activators and inhibitors found in this study provide valuable targets for further experiments to understand and control the function of *Vch*CAST.

Among the strong activators and inhibitors identified and validated within this study, we were particularly interested in the role of the RecBCD complex. RecD was identified as the strongest inhibitor of *Vch*CAST integration via the screen and Keio mutant validation (Figure 1B, 2A).

RecD functions as an inhibitor to homologous recombination by blocking RecA loading (33). Therefore it was unsurprising that RecA, a single-stranded binding protein essential for Rec-mediated double-strand break repair, showed an activating phenotype when single mutants were tested for *Vch*CAST integration efficiency (Figure 2B). Based on this low editing efficiency in the absence of *recA*, we’d expect decreased efficacy of *Vch*CAST in the Δ*recA* genotypes of many commercial *E. coli* strains. The weaker phenotypes of RecBC, may be explained by some amount of functional redundancy in RecBCD-mediated repair to other repair pathways, such as the RecF homologous recombination pathway (42). In the Mu transposon, the RecBCD complex facilitates the repair of a double-strand break that occurs during resolution of the insertion (43, 44). Similarly, we hypothesize that the highly stable *Vch*CAST post-transposition complex could stall replication forks and introduce a double-strand break at the target site, similar to Mu’s mechanism (6). RecBCD and RecA would then repair the double-strand break. This putative function is supported by our finding that the highly efficient homologous recombination system, λ-Red, improves the efficiency of *Vch*CAST. Such a homologous recombination-dependent mechanism for *Vch*CAST would contrast with the commonly stated assumption that transposition occurs independently of this DNA repair pathway (4, 6, 18, 45, 46). Further experiments would be necessary to validate this potential role of homologous recombination and RecBCD in *Vch*CAST function.

To examine the role of RecBCD and other effector proteins in *Vch*CAST function, we delivered *Vch*CAST via suicide vector. This approach leads to lower insertion efficiencies but provides a more dynamic range to examine the effects of activating and inhibiting proteins. Furthermore, using a suicide vector is advantageous for eventual microbiome editing applications since it is eliminated after cargo delivery, thus presenting fewer biocontainment concerns. The highest efficiencies reported in the literature are achieved in studies that deliver the *Vch*CAST on a replicating plasmid which is then selected for, where insertions can be introduced to nearly 100% of cells (8, 18, 45, 47). This is likely due to the presence of multiple copies of the *Vch*CAST vector persisting in cells for an extended time. Given that λ-Red has often been used to increase homologous recombination efficiency on a replicating vector (48), we expect that our λ-Red *Vch*CAST system would also work in that context.

Our findings on improving *Vch*CAST efficiency in *E. coli* prompted us to investigate the broader applicability of this approach in other bacterial species. When the λ-Red *Vch*CAST system was tested in Pseudomonadota outside of *E. coli*, we found that it enhanced editing efficiencies, but to a lesser extent (Figure 4A-B). These data suggest that insights from this screen will be most efficiently applied in a host-dependent manner. For example, commandeering the recombination machinery from a phage infecting the bacteria of interest and integrating it with the *Vch*CAST vector would likely allow for more effective editing.

Beyond Pseudomonadota, where type I-F CASTs are largely confined, many of the activators identified in this study have limited presence (Figure 5). Notably, IHF, which is essential for efficient *Vch*CAST transposition is absent from the majority of the bacterial tree of life. Given IHF’s importance in *E. coli*, it may be integral for targeting strains without endogenous IHF. For instance, members of Bascillota and Actinomycetota, where *Vch*CAST has shown to have low efficiency (7), lack close sequence homologs of IhfA and IhfB. While Actinomycetota does contain sequence-divergent but functionally homologous IHF-like proteins (49, 50), their ability to support transposition is unknown. The addition of *ihfA* and *ihfB* to *Vch*CAST editing vectors in these phyla, as well as other phyla where their homologs are absent, may increase editing efficiency. These considerations highlight the relevance of tailoring optimization strategies to the specific genetic characteristics of the target organism when applying *Vch*CAST systems more broadly.

**Figure 5:**
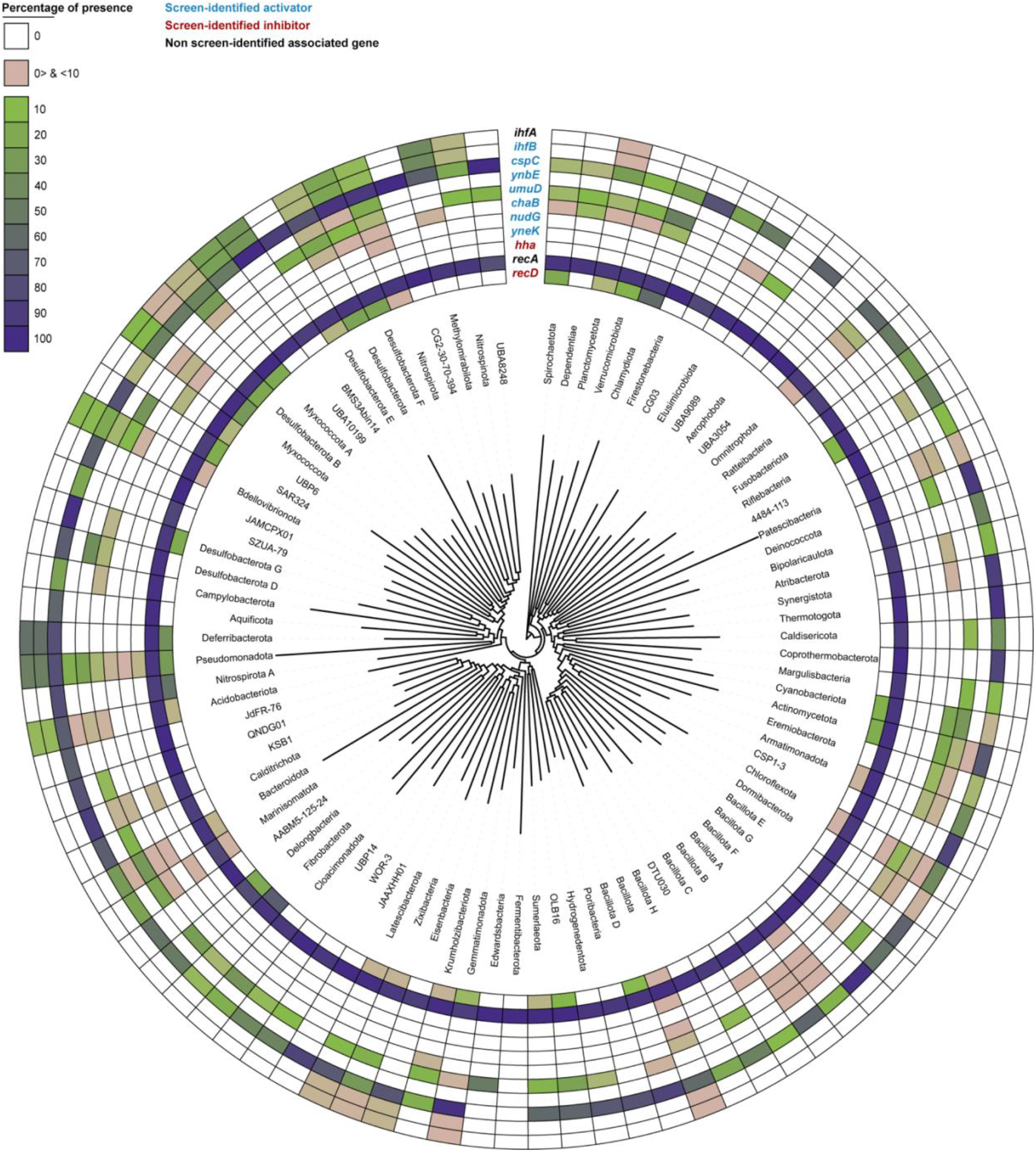
Phylogenetic distribution of *Vch*CAST activator and inhibitor genes. The phylogenetic distribution of 11 *E. coli* regulatory genes was mapped across 80,789 representative bacterial genomes from 92 phyla in Genome Taxonomy Database, release 214.0 (GTDB v214.0; (31, 32) with at least 10 members. Homologs were identified using AnnoTree (19) and confirmed with the eggNOG database (29).

We are excited about the potential of this work to bring the powerful CAST editing toolset to more biological systems. *Vch*CAST has mostly been applied in Gammaproteobacteria (5, 18, 51–55). Understanding of host factors that enable and inhibit these systems is an important barrier to their use across the phylogenetic tree of bacteria. In the complex microbiomes that are most relevant for human health and the environment, editing efficiency is a major bottleneck for delivering functional cargo insertions via *Vch*CAST (5). This work provides strategies for more efficient delivery so that the function of these communities can be better probed and controlled at a genetic level. Additionally, this work may inform the discovery of more activators for CAST systems in human cells, where efficiencies are currently low (6). Overall, our findings provide insights into the factors influencing CAST efficiency and offer strategies for improving its performance across diverse organisms, paving the way for broader applications in genome editing.

## DATA AVAILABILITY

Plasmids, strains, and oligonucleotides used in this study are available in the supplemental materials. RB-TnSeq barcode counts, fitness scores, and analysis scripts are available at https://figshare.com/account/home#/projects/220006. Custom scripts to analyze the Whole Genome Sequencing data and distribution of 11 genes in the bacterial tree of life are also at that link. Datasets generated and analyzed in the current study are available from the corresponding authors on reasonable request.

## SUPPLEMENTARY DATA

Supplementary Data are available at NAR online.

## ACKNOWLEDGEMENTS

We are immensely grateful to all of our collaborators for their contributions. We would like to thank Benjamin Adler, Jaymin Patel, Carlotta Ronda, and Robin Herbert for their input and discussion on the manuscript. We would also like to thank Morgan Price for his advice and support on RB-TnSeq analysis. We would like to extend our special thanks to the Arkin Lab for generously providing/supplying us with Keio collection mutants. We gratefully acknowledge the leadership and support of the Berkeley Initiative for Optimized Microbiome Editing (BIOME), particularly Jill Banfield, Brad Ringeisen, Audrey Glynn, and Rachel K. Evans. The schematics were generated by A.T.P.A using BioRender and are available under the following agreement numbers: LT279S5BCI, HP279S5ZMR, IY279S6DQQ, ZG279S6OMA.

## REFERENCES

1. Hsieh, S.-C. and Peters, J.E. (2024) Natural and Engineered Guide RNA-Directed Transposition with CRISPR-Associated Tn7-Like Transposons. Annu. Rev. Biochem., 93, 139–161.

2. Chang, C.-W., Truong, V.A., Pham, N.N. and Hu, Y.-C. (2024) RNA-guided genome engineering: paradigm shift towards transposons. Trends Biotechnol., 10.1016/j.tibtech.2024.02.006.

3. Vento, J.M., Crook, N. and Beisel, C.L. (2019) Barriers to genome editing with CRISPR in bacteria. J. Ind. Microbiol. Biotechnol., 46, 1327–1341.

4. Gelsinger, D.R., Vo, P.L.H., Klompe, S.E., Ronda, C., Wang, H.H. and Sternberg, S.H. (2024) Bacterial genome engineering using CRISPR-associated transposases. Nat. Protoc., 10.1038/s41596-023-00927-3.

5. Rubin, B.E., Diamond, S., Cress, B.F., Crits-Christoph, A., Lou, Y.C., Borges, A.L., Shivram, H., He, C., Xu, M., Zhou, Z., et al. (2022) Species- and site-specific genome editing in complex bacterial communities. Nat Microbiol, 7, 34–47.

6. Lampe, G.D., King, R.T., Halpin-Healy, T.S., Klompe, S.E., Hogan, M.I., Vo, P.L.H., Tang, S., Chavez, A. and Sternberg, S.H. (2024) Targeted DNA integration in human cells without double-strand breaks using CRISPR-associated transposases. Nat. Biotechnol., 42, 87–98.

7. Yang, S., Zhu, J., Zhou, X., Zhang, J., Li, Q., Bian, F., Zhu, J., Yan, T., Wang, X., Zhang, Y., et al. (2023) RNA-Guided DNA Transposition in Corynebacterium glutamicum and Bacillus subtilis. ACS Synth. Biol., 12, 2198–2202.

8. Walker, M.W.G., Klompe, S.E., Zhang, D.J. and Sternberg, S.H. (2023) Novel molecular requirements for CRISPR RNA-guided transposition. Nucleic Acids Res., 51, 4519–4535.

9. Wetmore, K.M., Price, M.N., Waters, R.J., Lamson, J.S., He, J., Hoover, C.A., Blow, M.J., Bristow, J., Butland, G., Arkin, A.P., et al. (2015) Rapid quantification of mutant fitness in diverse bacteria by sequencing randomly bar-coded transposons. MBio, 6, e00306–15.

10. Adler, B.A., Kazakov, A.E., Zhong, C., Liu, H., Kutter, E., Lui, L.M., Nielsen, T.N., Carion, H., Deutschbauer, A.M., Mutalik, V.K., et al. (2021) The genetic basis of phage susceptibility, cross-resistance and host-range in Salmonella. Microbiology, 167.

11. Wang, P., Yu, Z., Li, B., Cai, X., Zeng, Z., Chen, X. and Wang, X. (2015) Development of an efficient conjugation-based genetic manipulation system for Pseudoalteromonas. Microb. Cell Fact., 14, 11.

12. Baba, T., Ara, T., Hasegawa, M., Takai, Y., Okumura, Y., Baba, M., Datsenko, K.A., Tomita, M., Wanner, B.L. and Mori, H. (2006) Construction of Escherichia coli K-12 in-frame, single-gene knockout mutants: the Keio collection. Mol. Syst. Biol., 2, 2006.0008.

13. Datsenko, K.A. and Wanner, B.L. (2000) One-step inactivation of chromosomal genes in Escherichia coli K-12 using PCR products. Proc. Natl. Acad. Sci. U. S. A., 97, 6640–6645.

14. Egbert, R.G., Rishi, H.S., Adler, B.A., McCormick, D.M., Toro, E., Gill, R.T. and Arkin, A.P. (2019) A versatile platform strain for high-fidelity multiplex genome editing. Nucleic Acids Res., 47, 3244–3256.

15. Ruegg, T.L., Pereira, J.H., Chen, J.C., DeGiovanni, A., Novichkov, P., Mutalik, V.K., Tomaleri, G.P., Singer, S.W., Hillson, N.J., Simmons, B.A., et al. (2018) Jungle Express is a versatile repressor system for tight transcriptional control. Nat. Commun., 9, 3617.

16. De Coster, W., D’Hert, S., Schultz, D.T., Cruts, M. and Van Broeckhoven, C. (2018) NanoPack: visualizing and processing long-read sequencing data. Bioinformatics, 34, 2666–2669.

17. Li, H. (2018) Minimap2: pairwise alignment for nucleotide sequences. Bioinformatics, 34, 3094–3100.

18. Vo, P.L.H., Ronda, C., Klompe, S.E., Chen, E.E., Acree, C., Wang, H.H. and Sternberg, S.H. (2021) CRISPR RNA-guided integrases for high-efficiency, multiplexed bacterial genome engineering. Nat. Biotechnol., 39, 480–489.

19. Mendler, K., Chen, H., Parks, D.H., Lobb, B., Hug, L.A. and Doxey, A.C. (2019) AnnoTree: visualization and exploration of a functionally annotated microbial tree of life. Nucleic Acids Res., 47, 4442–4448.

20. Finn, R.D., Coggill, P., Eberhardt, R.Y., Eddy, S.R., Mistry, J., Mitchell, A.L., Potter, S.C., Punta, M., Qureshi, M., Sangrador-Vegas, A., et al. (2016) The Pfam protein families database: towards a more sustainable future. Nucleic Acids Res., 44, D279–85.

21. Haft, D.H., Selengut, J.D. and White, O. (2003) The TIGRFAMs database of protein families. Nucleic Acids Res., 31, 371–373.

22. Suzek, B.E., Wang, Y., Huang, H., McGarvey, P.B., Wu, C.H. and UniProt Consortium (2015) UniRef clusters: a comprehensive and scalable alternative for improving sequence similarity searches. Bioinformatics, 31, 926–932.

23. Parks, D.H., Chuvochina, M., Rinke, C., Mussig, A.J., Chaumeil, P.-A. and Hugenholtz, P. (2022) GTDB: an ongoing census of bacterial and archaeal diversity through a phylogenetically consistent, rank normalized and complete genome-based taxonomy. Nucleic Acids Res., 50, D785–D794.

24. Haluzi, H., Goitein, D., Koby, S., Mendelson, I., Teff, D., Mengeritsky, G., Giladi, H. and Oppenheim, A.B. (1991) Genes coding for integration host factor are conserved in gram-negative bacteria. J. Bacteriol., 173, 6297–6299.

25. Sutton, M.D., Opperman, T. and Walker, G.C. (1999) The Escherichia coli SOS mutagenesis proteins UmuD and UmuD’ interact physically with the replicative DNA polymerase. Proc. Natl. Acad. Sci. U. S. A., 96, 12373–12378.

26. Bernheim, A., Bikard, D., Touchon, M. and Rocha, E.P.C. (2019) A matter of background: DNA repair pathways as a possible cause for the sparse distribution of CRISPR-Cas systems in bacteria. Philos. Trans. R. Soc. Lond. B Biol. Sci., 374, 20180088.

27. Graumann, P.L. and Marahiel, M.A. (1998) A superfamily of proteins that contain the cold-shock domain. Trends Biochem. Sci., 23, 286–290.

28. Catalan-Moreno, A., Caballero, C.J., Irurzun, N., Cuesta, S., López-Sagaseta, J. and Toledo-Arana, A. (2020) One evolutionarily selected amino acid variation is sufficient to provide functional specificity in the cold shock protein paralogs of Staphylococcus aureus. Mol. Microbiol., 113, 826–840.

29. Huerta-Cepas, J., Szklarczyk, D., Heller, D., Hernández-Plaza, A., Forslund, S.K., Cook, H., Mende, D.R., Letunic, I., Rattei, T., Jensen, L.J., et al. (2019) eggNOG 5.0: a hierarchical, functionally and phylogenetically annotated orthology resource based on 5090 organisms and 2502 viruses. Nucleic Acids Res., 47, D309–D314.

30. Letunic, I. and Bork, P. (2024) Interactive Tree of Life (iTOL) v6: recent updates to the phylogenetic tree display and annotation tool. Nucleic Acids Res., 52, W78–W82.

31. Parks, D.H., Chuvochina, M., Waite, D.W., Rinke, C., Skarshewski, A., Chaumeil, P.-A. and Hugenholtz, P. (2018) A standardized bacterial taxonomy based on genome phylogeny substantially revises the tree of life. Nat. Biotechnol., 36, 996–1004.

32. Parks, D.H., Chuvochina, M., Chaumeil, P.-A., Rinke, C., Mussig, A.J. and Hugenholtz, P. (2020) A complete domain-to-species taxonomy for Bacteria and Archaea. Nat. Biotechnol., 38, 1079–1086.

33. Amundsen, S.K., Taylor, A.F. and Smith, G.R. (2000) The RecD subunit of the Escherichia coli RecBCD enzyme inhibits RecA loading, homologous recombination, and DNA repair. Proc. Natl. Acad. Sci. U. S. A., 97, 7399–7404.

34. Vo, P.L.H., Acree, C., Smith, M.L. and Sternberg, S.H. (2021) Unbiased profiling of CRISPR RNA-guided transposition products by long-read sequencing. Mob. DNA, 12, 13.

35. Murphy, K.C. (2007) The lambda Gam protein inhibits RecBCD binding to dsDNA ends. J. Mol. Biol., 371, 19–24.

36. Mosberg, J.A., Lajoie, M.J. and Church, G.M. (2010) Lambda red recombineering in Escherichia coli occurs through a fully single-stranded intermediate. Genetics, 186, 791– 799.

37. Weimer, A., Kohlstedt, M., Volke, D.C., Nikel, P.I. and Wittmann, C. (2020) Industrial biotechnology of Pseudomonas putida: advances and prospects. Appl. Microbiol. Biotechnol., 104, 7745–7766.

38. Yu, Z., Li, S., Li, Y., Jiang, Z., Zhou, J. and An, Q. (2018) Complete genome sequence of N2-fixing model strain Klebsiella sp. nov. M5al, which produces plant cell wall-degrading enzymes and siderophores. Biotechnol Rep (Amst), 17, 6–9.

39. Prah, I., Nukui, Y., Yamaoka, S. and Saito, R. (2022) Emergence of a High-Risk Klebsiella michiganensis Clone Disseminating Carbapenemase Genes. Front. Microbiol., 13, 880248.

40. Chaurasiya, K.R., Ruslie, C., Silva, M.C., Voortman, L., Nevin, P., Lone, S., Beuning, P.J. and Williams, M.C. (2013) Polymerase manager protein UmuD directly regulates Escherichia coli DNA polymerase III α binding to ssDNA. Nucleic Acids Res., 41, 8959–8968.

41. Phadtare, S. and Inouye, M. (1999) Sequence-selective interactions with RNA by CspB, CspC and CspE, members of the CspA family of Escherichia coli. Mol. Microbiol., 33, 1004–1014.

42. Dillingham, M.S. and Kowalczykowski, S.C. (2008) RecBCD enzyme and the repair of double-stranded DNA breaks. Microbiol. Mol. Biol. Rev., 72, 642–71, Table of Contents.

43. Choi, W., Jang, S. and Harshey, R.M. (2014) Mu transpososome and RecBCD nuclease collaborate in the repair of simple Mu insertions. Proc. Natl. Acad. Sci. U. S. A., 111, 14112–14117.

44. Jang, S., Sandler, S.J. and Harshey, R.M. (2012) Mu insertions are repaired by the double-strand break repair pathway of Escherichia coli. PLoS Genet., 8, e1002642.

45. Klompe, S.E., Vo, P.L.H., Halpin-Healy, T.S. and Sternberg, S.H. (2019) Transposon-encoded CRISPR–Cas systems direct RNA-guided DNA integration. Nature, 571, 219–225.

46. Wei, J. and Li, Y. (2023) CRISPR-based gene editing technology and its application in microbial engineering. Engineering Microbiology, 3, 100101.

47. Roberts, A., Nethery, M.A. and Barrangou, R. (2022) Functional characterization of diverse type I-F CRISPR-associated transposons. Nucleic Acids Res., 50, 11670–11681.

48. Fels, U., Gevaert, K. and Van Damme, P. (2020) Bacterial Genetic Engineering by Means of Recombineering for Reverse Genetics. Front. Microbiol., 11, 548410.

49. Sharadamma, N., Harshavardhana, Y., Ravishankar, A., Anand, P., Chandra, N. and Muniyappa, K. (2014) Molecular dissection of Mycobacterium tuberculosis integration host factor reveals novel insights into the mode of DNA binding and nucleoid compaction. J. Biol. Chem., 289, 34325–34340.

50. Swiercz, J.P., Nanji, T., Gloyd, M., Guarné, A. and Elliot, M.A. (2013) A novel nucleoid-associated protein specific to the actinobacteria. Nucleic Acids Res., 41, 4171–4184.

51. Zhang, Y., Sun, X., Wang, Q., Xu, J., Dong, F., Yang, S., Yang, J., Zhang, Z., Qian, Y., Chen, J., et al. (2020) Multicopy Chromosomal Integration Using CRISPR-Associated Transposases. ACS Synth. Biol., 9, 1998–2008.

52. Yang, S., Zhang, Y., Xu, J., Zhang, J., Zhang, J., Yang, J., Jiang, Y. and Yang, S. (2021) Orthogonal CRISPR-associated transposases for parallel and multiplexed chromosomal integration. Nucleic Acids Res., 49, 10192–10202.

53. Banta, A.B., Myers, K.S., Ward, R.D., Cuellar, R.A., Place, M., Freeh, C.C., Bacon, E.E. and Peters, J.M. (2024) A Targeted Genome-scale Overexpression Platform for Proteobacteria. bioRxiv, 10.1101/2024.03.01.582922.

54. Xu, J., Sun, Y., Wu, J., Yang, S. and Yang, L. (2024) Chromosome recombination and modification by LoxP-mediated evolution in Vibrio natriegens using CRISPR-associated transposases. Biotechnol. Bioeng., 121, 1163–1172.

55. Garza Elizondo, A.M. and Chappell, J. (2024) Targeted Transcriptional Activation Using a CRISPR-Associated Transposon System. ACS Synth. Biol., 13, 328–336.

