## Supplementary Figures for "Identification of Proteins Influencing CRISPR-Associated Transposases for Enhanced Genome Editing"

**MANUSCRIPT TITLE**

**SUPPLEMENTAL FIGURES**

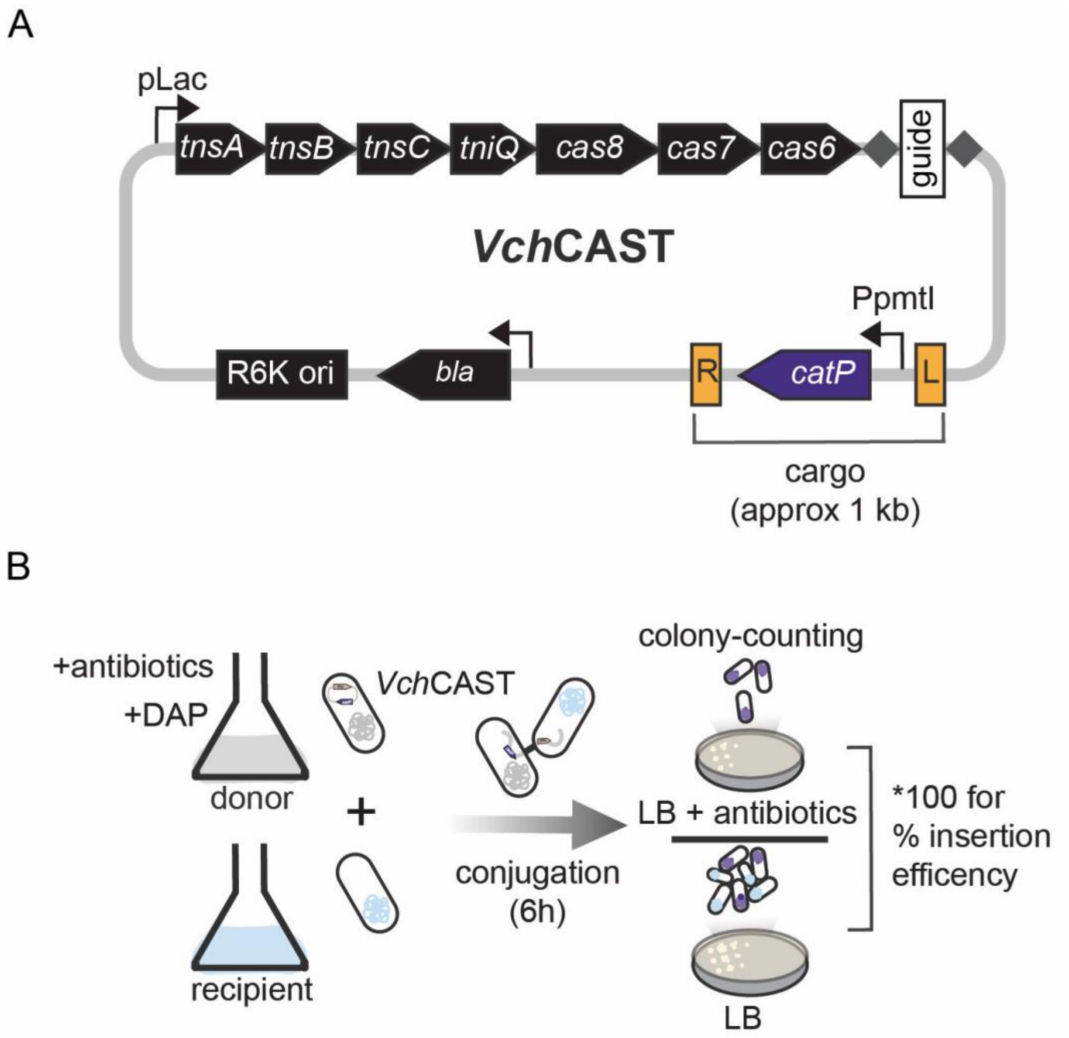


**Figure S1.** *VchCAST construct and conjugative delivery-insertion efficiency assay.* (A) Plasmid schematic for the standard type I-F *Vch*CAST editing vector. (B) Conceptual schematic for conjugation-based *Vch*CAST insertion efficiency assay.

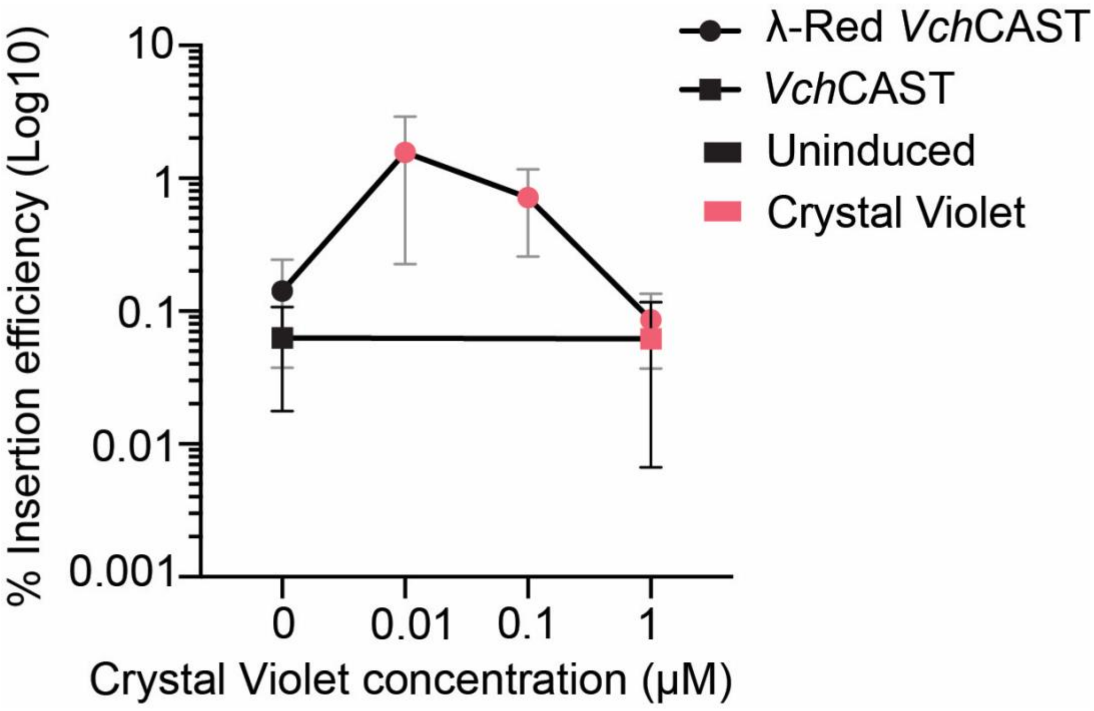


**Figure S2.** *CV induction optimization for λ-Red expression in E. coli.*

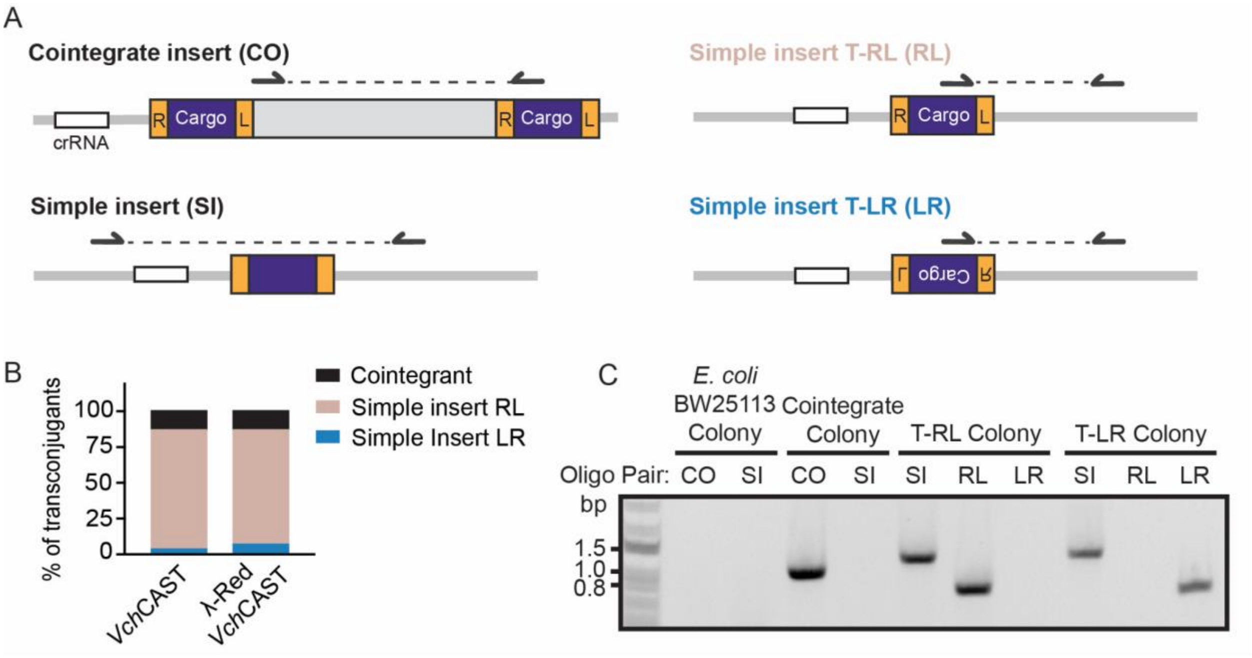


**Figure S3.** *Comparison of insertion products for VchCAST and λ-Red VchCAST in BW25113 E. coli.* (A) Schematic of primer binding and amplification for insertion product verification and orientation analysis by colony PCR (cPCR). (B) Side-by-side comparison of product type and orientation (%) of *Vch*CAST and λ-Red *Vch*CAST in *E. coli* BW25113 as characterized by cPCR. (C) Representative gel image of cPCR analysis of *Vch*CAST insertion products in *E. coli* BW25113. Oligo pairs used for amplification are detailed in part A of this figure. CO is cointegrate, SI is simple insert, and the orientations are denoted as RL (right-left) and LR (left-right).

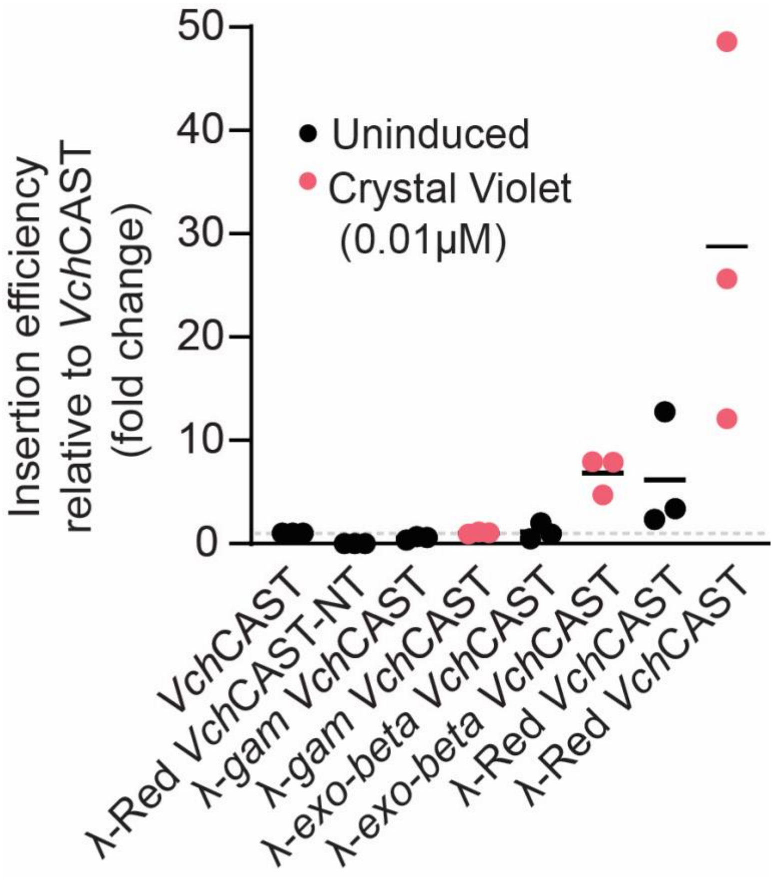


**Figure S4.** *Relative editing efficiency of VchCAST vectors containing different combinations of λ-Red genes.* The editing efficiency of λ-Red *Vch*CAST as well as the non-targeting (NT) control were normalized to paired *Vch*CAST biological replicates.

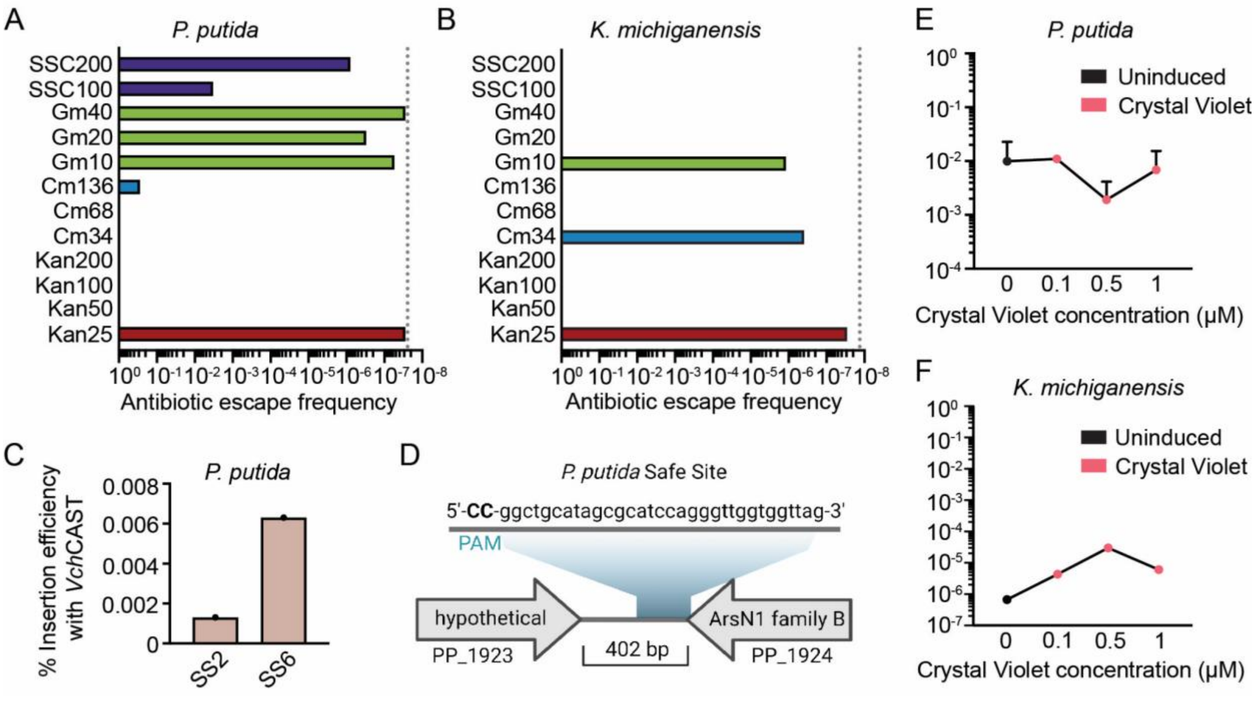


**Figure S5.** *P. Putida and K. michiganensis VchCAST editing.* (A) Antibiotic susceptibility testing for *P. putida*. The dotted line depicts the limit of detection 2.56E$^{-8}$ (B) Antibiotic susceptibility testing for *K. michiganensis*. The dotted line depicts the limit of detection 1.33E$^{-8}$ (C) Guide design for *P. putida* Safe site 6 (SS6). (D) Safe site guide insertion efficiency (%) in *P. putida*. Created in BioRender. Alker, A. (2024) BioRender.com/m51i476 (E) CV induction optimization for λ-Red expression in *P. putida*. (F) CV induction optimization for λ-Red expression in *K. michiganensis*.
